# CNPY4 inhibits the Hedgehog pathway by modulating membrane sterol lipids

**DOI:** 10.1101/2021.03.15.435490

**Authors:** Megan Lo, Amnon Sharir, Michael D. Paul, Hayarpi Torosyan, Christopher Agnew, David R. Raleigh, Natalia Jura, Ophir D. Klein

**Author notes:** Correspondence should be addressed to N.J. or O.D.K.

## Abstract

The Hedgehog (HH) pathway is critical for development and adult tissue homeostasis^1^. Aberrant HH signaling can cause congenital malformations, such as digit anomalies and holoprosencephaly^2^, and other diseases, including cancer^3^. Signal transduction is initiated by HH ligand binding to the Patched 1 (PTCH1) receptor on primary cilia, thereby releasing inhibition of Smoothened (SMO), a HH pathway activator^4^. Although cholesterol and several oxysterol lipids, which are enriched in the ciliary membrane, play a crucial role in HH activation^4,5^, the molecular mechanisms governing the regulation of these lipid molecules remain unresolved. Here, we identify Canopy 4 (CNPY4), a Saposin-like protein, as a regulator of the HH pathway that controls membrane sterol lipid levels. *Cnpy4*^−/−^ embryos exhibit multiple defects consistent with HH signaling perturbations, most notably changes in digit number. Knockdown of *Cnpy4* hyperactivates the HH pathway at the level of SMO *in vitro*, and elevates membrane levels of accessible sterol lipids such as cholesterol, an endogenous ligand involved in SMO activation^6^. Thus, our data demonstrate that CNPY4 is a negative regulator that fine-tunes the initial steps of HH signal transduction, revealing a previously undescribed facet of HH pathway regulation that operates through control of membrane composition.

## Main body

The hedgehog (*HH*) gene was first identified in *Drosophila* as a regulator of larval segmentation^7^, after which three mammalian homologs were discovered: desert hedgehog (*Dhh*), Indian hedgehog (*Ihh*), and sonic hedgehog (*Shh*)^8–11^. *Shh* is the most widely expressed HH ligand and is found in the epithelium and at epithelial-mesenchymal boundaries of various tissues, including the tooth, gut, lung, and limb, where it controls morphogenesis and adult homeostasis^12^. Precise regulation of *Shh* signaling is therefore critical for proper tissue development and patterning. Perturbations to the pathway have been linked to severe congenital abnormalities, including polydactyly and holoprosencephaly^12^. Misregulation of *Shh* pathway genes can also lead to cancers such as basal cell carcinoma, the most common cancer in the United States, and medulloblastoma, the most common malignant brain cancer in children^2,3^.

HH signal transduction in vertebrates occurs through a tightly regulated process at the primary cilium, an antenna-like organelle that protrudes from the surface of most cells^13,14^. Signaling is initiated by binding of a secreted HH ligand to the PTCH1 receptor, which resides in and at the base of primary cilia^15–19^. HH binding to PTCH1 releases inhibition of the G-protein coupled receptor SMO, leading to SMO accumulation in cilia^20^. There, SMO is likely activated by one or more sterol lipid ligands^21^, whose exact identities remain to be unequivocally determined^22^. Activation of SMO releases the inhibition of the glioma-associated oncogene (GLI) transcription factors (GLI1, 2, and 3) by a negative regulator of the pathway, Suppressor of Fused (SUFU)^23^. This allows the GLI proteins to translocate into the nucleus and initiate transcription of key developmental genes^24–26^. HH activation also upregulates transcription of pathway genes including *Ptch1* and *Gli1*, leading to a complex signaling feedback loop^27,28^. Additionally, several components of the HH pathway interact with sterol lipids, and both depletion of cellular lipids and inhibition of sterol biosynthesis hinders HH signal transduction^29–35^. These lipid molecules are thought to influence trafficking of proteins into and out of, and thereby signal transduction from, the cilia^5^. Thus, sterol localization and concentration at the plasma membrane is critically tied to HH signaling.

We set out to explore the potential role of the Canopy (CNPY) subfamily of saposin and saposin-like (SAPLIP) proteins in the regulation of HH signaling. A number of SAPLIP proteins interact with lipids to modulate processes such as membrane binding, permeabilization, and lipid metabolism. In zebrafish, *cnpy1* was reported to regulate the development of Kupffer’s vesicle, which controls left-right asymmetry through HH signaling in zebrafish^36–38^; however, the interaction of *cnpy1* with HH signaling was not explored. In humans and mice, *Cnpy1* appears truncated, and *Cnpy2* and *Cnpy3* knockout mice do not display visible alterations of HH pathway^36,39–41^. We therefore turned to the remaining family member, *Cnpy4*, to study whether it may play a role in HH signaling. We bred *Cnpy4* knockout mouse lines and assessed the effect of CNPY4 loss of function in mutant embryos (Fig. 1; Extended Data Fig. 1). Of the *Cnpy4*^−/−^ embryos examined, 85% exhibited abnormalities in hindlimb digit number, ranging from the formation of one or two supernumerary digits on the anterior side of the limb (termed preaxial polydactyly) to a loss of up to three posterior digits (Fig. 1c, Extended Data Fig. 1a, b). Similar bi-directional phenotypic changes to the limb buds have also been observed in patients with loss of function *Gli3* mutations^42,43^. Approximately 20% of the *Cnpy4*^−/−^ mutants exhibited other anomalies consistent with HH pathway misregulation, including rostral and/or caudal neural tube closure defects, splayed vertebrae, and abnormal rib morphology with fusions and bifurcations^44,45^ (Extended Data Fig. 1c). Due to the high penetrance of the limb phenotype and the central role of *Shh* in controlling digit number, we focused further analysis on limb abnormalities in *Cnpy4* knockout mice.

**Fig. 1.**
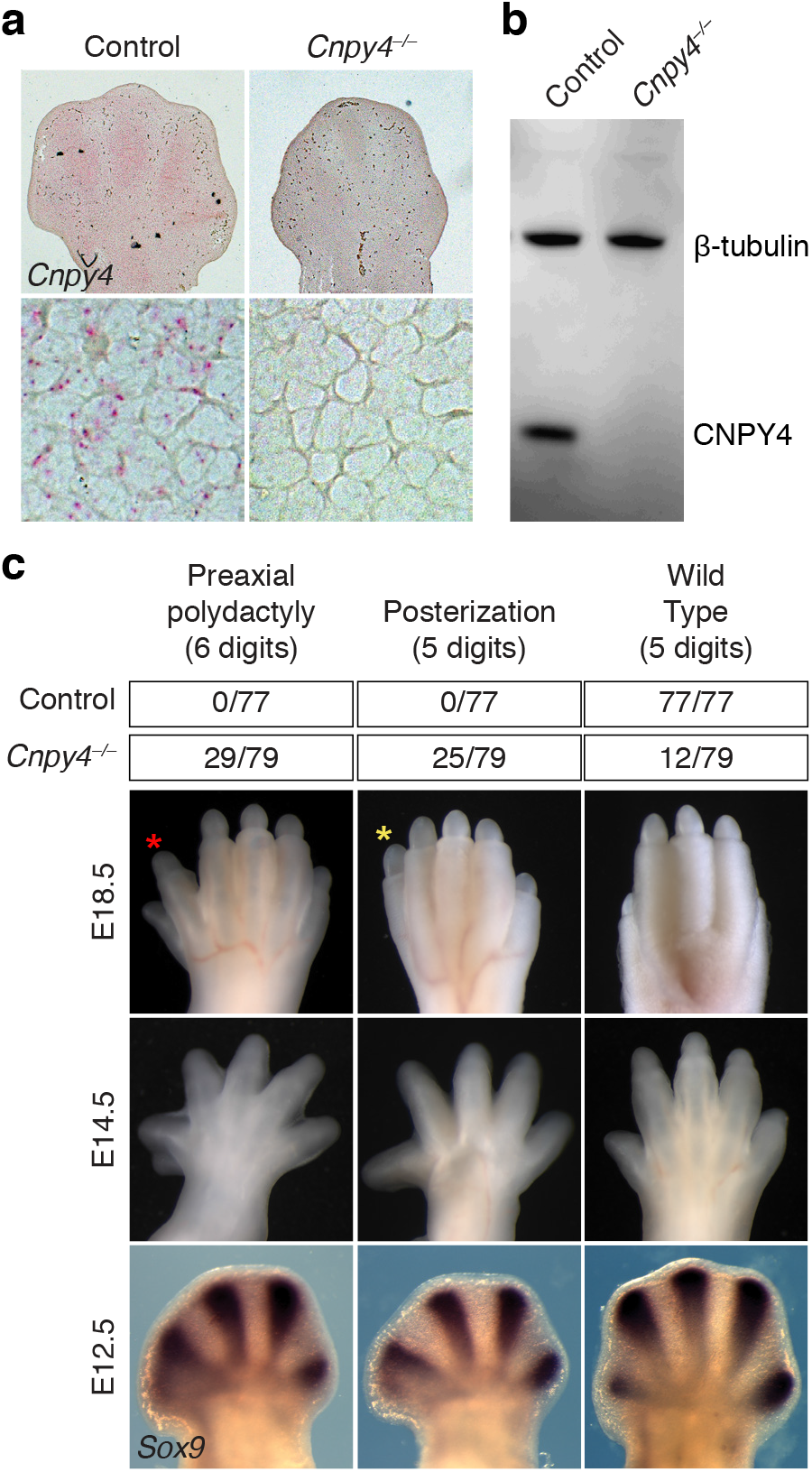
Developmental defects in *Cnpy4*^−/−^ hindlimbs. **a**, Single-molecule *in situ* hybridization (RNAscope) of *Cnpy4* in hindlimbs of wild-type and *Cnpy4* mutant embryos at E12.5. Boxed areas are magnified on the bottom. **b**, Protein extracts of MEFs from control and mutant embryos were blotted with an anti-CNPY4 antibody and an β-tubulin loading control antibody. **c**, Dorsal view of wild-type and *Cnpy4* mutant limbs at E18.5 (top row) and E14.5 (middle row). The majority of *Cnpy4* hindlimbs exhibit either an extra digit anteriorly (yellow asterisk) or a transformation of digit 1 from biphalangeal to triphalangeal (red asterisk). The top table summarizes the phenotype frequency in mutant hindlimbs; less frequent phenotypes are shown in Extended Figure 1a, b Whole mount in situ hybridization for *Sox9* (bottom row) indicates an extra digit (arrow) and an enlarged digit 1 primordium.

To explore whether *Cnpy4* modulates *Shh*, we first examined the expression of *Shh* and its downstream effector *Gli1* during limb development in mutant embryos. *Shh* and *Gli1* expression expanded anteriorly in the early hindlimb buds of *Cnpy4* mutants (embryonic day (E) 10.5 - E11.5), and ectopic expression of *Shh* and *Gli1* was present in anterior domains at later developmental stages (E12.5) (Fig. 2a), in line with misactivation of the HH pathway. These changes are consistent with those observed in other human patients and mouse models with preaxial polydactyly^46–48^. In a small number of mutants, reduction of *Shh* and *Gli1* expression was observed (Extended Data Fig. 2), paralleling the minority of *Cnpy4*^−/−^ mutants manifesting oligodactyly.

**Fig. 2.**
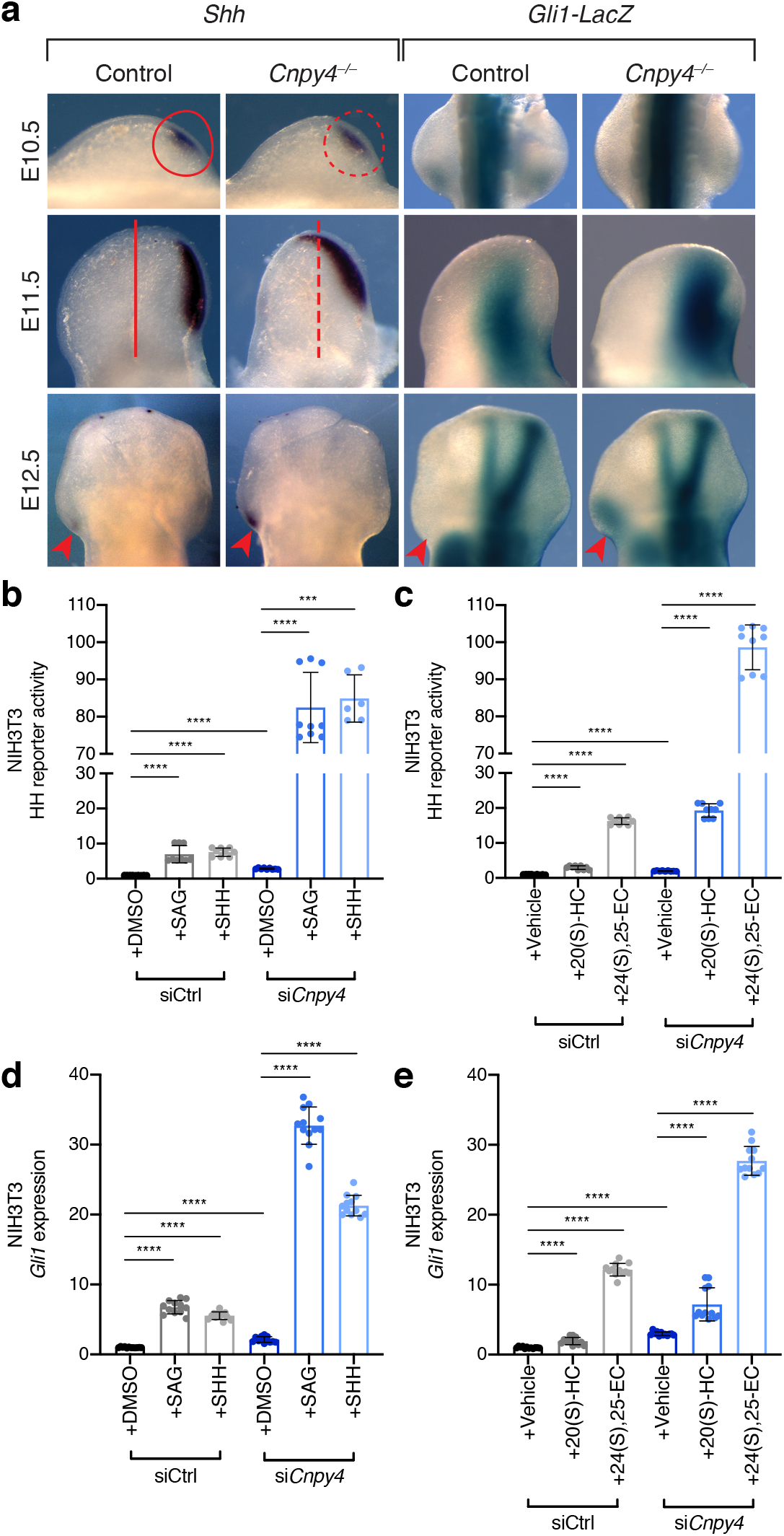
Absence of *Cnpy4* leads to hyperactivation of HH-related gene expression and signaling. **a**, *In situ* hybridization and lacZ expression of *Shh* and *Gli1-lacZ* in hindlimb buds at E10.5, E11.5 and E12.5 showing enlarged *Shh* domain (circles and lines) and ectopic expression of both *Shh* and *Gli1* (arrrowheads) in the *Cnpy4* mutants. **b, c**, Luciferase reporter assay in ciliated NIH3T3 cells treated with *Cnpy4* or control siRNA and stimulated with SAG or recombinant SHH (**b**) and 20(S)-hydroxycholesterol or 24(S), 25-epoxycholesterol (**c**). Quantifications were normalized to the average value of control siRNA treated cells stimulated with DMSO or vehicle. **d, e**, qRT-PCR assessment of *Gli1* expression in ciliated NIH3T3 cells treated with *Cnpy4* or control siRNA and stimulated with SAG or recombinant SHH (**d**) and 20(S)-hydroxycholesterol or 24(S), 25-epoxycholesterol (**e**). Significance calculations were performed as described in Methods and Materials; **** p < 0.0001

In order to measure the *Cnpy4*-dependent changes in HH signaling at the cellular level, we utilized a luciferase reporter assay to measure *Gli* expression in NIH3T3 cells following transient *Cnpy4* knockdown with siRNA (Extended Data Fig. 3a). Consistent with the predominant polydactyly phenotype and other developmental abnormalities we observed in *Cnpy4* knockout embryos, silencing of *Cnpy4* resulted in elevated basal activation of the HH transcriptional program and potentiated signaling in response to HH pathway agonists (Fig. 2b, c). These effects were independent of the ligand used to activate the pathway, including a chemical SMO agonist (SAG), recombinant SHH, and both synthetic (20(S)-hydroxycholesterol) and cilia-associated (24(S), 25-epoxycholesterol) oxysterols that bind and activate SMO^44^. To corroborate these results, we directly analyzed *Gli1* transcript levels in NIH3T3 cells using qRT-PCR. In line with the results from the HH luciferase reporter assay, we found that *Gli1* expression was greatly increased in *Cnpy4* knockdown cells compared to those treated with control siRNA upon ligand stimulation (Fig. 2d, e; Extended Data Fig. 3b, c). Thus, the HH pathway is hyperactive in cells lacking *Cnpy4*, suggesting that CNPY4 is a negative regulator of the HH pathway.

Morphological differences in cilia, changes in cell ciliation, and improper trafficking of ciliary proteins are all linked to aberrant HH activity during development^5,13,14,49–51^. We therefore asked if ciliary defects could explain the hyperactivation of the HH pathway observed by staining for acetylated tubulin, a marker of the ciliary axoneme, in *Cnpy4* deficient NIH3T3 cells and in mouse embryonic fibroblasts (MEFs) derived from *Cnpy4*^−/−^ embryonic limb buds (Fig. 3a; Extended Data Fig. 4a–c). NIH3T3 cells with *Cnpy4* knockdown and *Cnpy4*^−/−^ MEFs did not show significant differences in the percentage of ciliated cells compared to control cells (Fig. 3b; Extended Data Fig. 4d). Furthermore, the length and overall morphology of cilia were not visibly altered by depletion of CNPY4 (Fig. 3a, c; Extended Data Fig. 4c, e). The intensity of SMO staining in the cilia upon SAG stimulation was also unchanged in *Cnpy4* silenced cells, indicating that the ability of SMO to traffic into the cilia was not impaired (Extended Data Fig. 4f, g). Similar uncoupling of ciliary morphology and SMO trafficking from HH activation were recently reported upon ablation of the cholesterol biosynthesis enzyme DHCR7^52^. Thus, we concluded that the effect CNPY4 exerts on the HH pathway was likely through signaling-specific events, rather than ciliary or protein compartmentalization abnormalities.

**Fig. 3.**
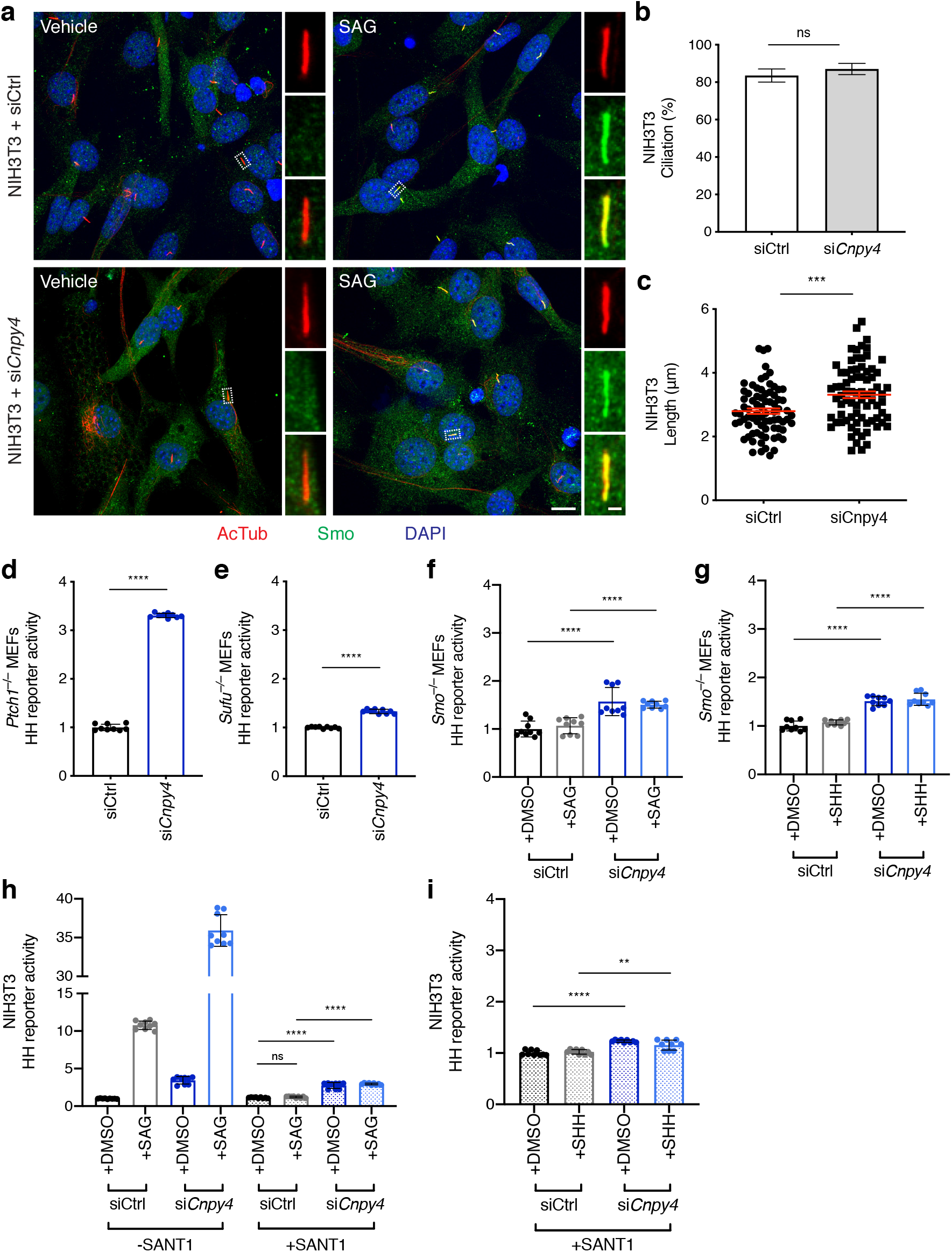
CNPY4 intersects the HH pathway at the level of SMO. **a**, Immunofluorescence of primary cilia (acetylated tubulin, red), SMO (SMO, green), and nuclei (DAPI, blue) in ciliated NIH3T3 cells treated with *Cnpy4* or control siRNA. The scale bar represents 10 um. Inset scale bar represents 1 um. **b**, Quantification of number of NIH3T3 cells ciliated as assessed by acetylated tubulin immunofluorescence. **c**, Quantification of ciliary length in NIH3T3 cells. Measurements were performed in FIJI using the acetylated tubulin channel. **d–g**, Luciferase reporter assay in ciliated *Ptch1* (**d**), *Sufu* (**e**), or *Smo* (**f, g**) null MEFs treated with *Cnpy4* or control siRNA, with *Smo* null MEFs stimulated with either SAG (**f**) or SHH (**g**). Quantifications were normalized to the average value of control siRNA treated cells. **h, i**, Luciferase reporter assay in ciliated NIH3T3 cells treated with *Cnpy4* or control siRNA stimulated with SAG (**h**) or SHH (**i**) in the presence of SANT-1. Quantifications were normalized to the average value of control siRNA treated cells stimulated with DMSO. All significance calculations were performed as described in Methods and Materials; ** p <0.005, **** p < 0.0001.

To map the impact CNPY4 exerts on HH signal transduction components, we utilized our *in vitro* system to perform epistasis experiments. Although knockout of *Ptch1* alone constitutively activates the HH pathway^20^, knockdown of *Cnpy4* further activated the HH transcriptional program in *Ptch1*^−/−^ MEFs compared to control cells (Fig. 3d; Extended Data Fig. 5a–c), suggesting CNPY4 intersects the HH pathway parallel to or downstream of PTCH1. Knockout of *Sufu*, a negative regulator of the pathway downstream of PTCH1, also results in constitutive activation of the HH pathway^53^ (Fig. 3e; Extended Data Fig. 5d–f). However, in contrast to the effect of *Cnpy4* knockdown in *Ptch1*^−/−^ MEFs, knockdown of *Cnpy4* in *Sufu*^−/−^ MEFs resulted in a comparatively modest increase of *Gli1* mRNA transcription, suggesting CNPY4 functions upstream of SUFU to inhibit HH signal transduction.

SMO functions downstream from PTCH1 and upstream from SUFU^9^. Since *Smo^−/−^* MEFs are unable to transduce HH signals in response to pathway ligands, we examined whether the observed CNPY4-mediated modulation of HH signaling required SMO using both genetic (Fig. 3f, g; Extended Data Fig. 6a, b) and pharmacological (Fig. 3h, i; Extended Data Fig. 6c–e) perturbations. Remarkably, in the absence of SMO, SAG or recombinant SHH stimulation was unable to elicit hyperactive HH signaling after *Cnpy4* knockdown (Fig. 3f, g; Extended Data Fig. 6b), indicating that, like PTCH1, CNPY4 modulates HH activity through SMO. This lack of hyperactivation was also observed in *Cnpy4*-silenced NIH3T3 cells when SMO was pharmacologically inhibited by its antagonist SANT-1, which directly competes with SAG for binding to SMO (Fig. 3h, i; Extended Data Fig. 6d). We noted that these cells displayed slightly elevated levels of basal HH activity upon knockdown of *Cnpy4*, despite the absence or repression of SMO in these cells (Fig. 3f–i; Extended Data Fig. 6b, d), although to a much lesser extent than cells expressing SMO. Together these findings point to an essential role of SMO in the ligand-dependent potentiating effect of CNPY4 loss on HH signaling.

As SMO and PTCH1 are both transmembrane proteins whose signaling is likely sensitive to the local lipid environment, we asked if CNPY4, as a SAPLIP protein, may modulate the lipid composition of the membrane. In comparison to other membrane compartments, the ciliary membrane in which PTCH1 and SMO reside is highly enriched in cholesterol and oxysterols^54^. These lipids have been shown to bind and activate SMO^21,55,56^. We therefore probed the ability of CNPY4 to interact with cholesterol and several of these oxysterol compounds *in vitro.* We purified a recombinant construct of human CNPY4 (CNPY4ΔCt) lacking its signal sequence and C-terminal tail, which is predicted to be largely unstructured (Extended Data Fig. 7a). Purified CNPY4ΔCt is well-folded and predominantly alpha helical, as expected for a SAPLIP protein (Extended Data Fig. 7b). However, under the conditions tested, recombinant CNPY4ΔCt did not appear to bind cholesterol (Extended Data Fig. 7c, d). Furthermore, purified CNPY4ΔCt did not display measurable binding to a number of oxysterols known to be specifically enriched in the ciliary membrane and directly involved in HH pathway activation^44^ (Extended Data Fig. 7e). As the ability of many SAPLIP proteins to interact with lipids is directly tied to their dimerization^57–61^, we tested if CNPY4 is a dimer. Size exclusion chromatography of recombinant CNPY4ΔCt (Extended Data Fig. 7a) and co-immunoprecipitation between two differentially tagged constructs of full-length CNPY4 (Extended Data Fig. 7e, f) are consistent with CNPY4 being a monomer. Thus, CNPY4 likely modulates the lipid membrane beyond direct interaction with its lipid components.

We therefore tested the possibility that the absence of CNPY4 could increase the membrane levels of unbound accessible sterols, among them cholesterol^62^, which is most abundant in animal plasma membranes and was recently proposed to be a ligand responsible for SMO activation^6^. To directly measure the levels of accessible sterols in the plasma membrane of intact cells, we used a modified protein probe derived from the bacterial toxin Perfringolysin O (PFO*) coupled to a fluorescent tag^62^. Remarkably, NIH3T3 cells in which *Cnpy4* was knocked down displayed significantly elevated levels of accessible sterols compared to control treated cells (Fig. 4a, b). Additionally, MEFs derived from embryonic limb buds of *Cnpy4* null animals had notably increased levels of accessible sterols in a basal state (Fig. 4c). These data indicate that the inhibitory effect of CNPY4 on the HH pathway is likely a consequence of decreased levels of sterol lipids at the plasma membrane.

**Fig. 4.**
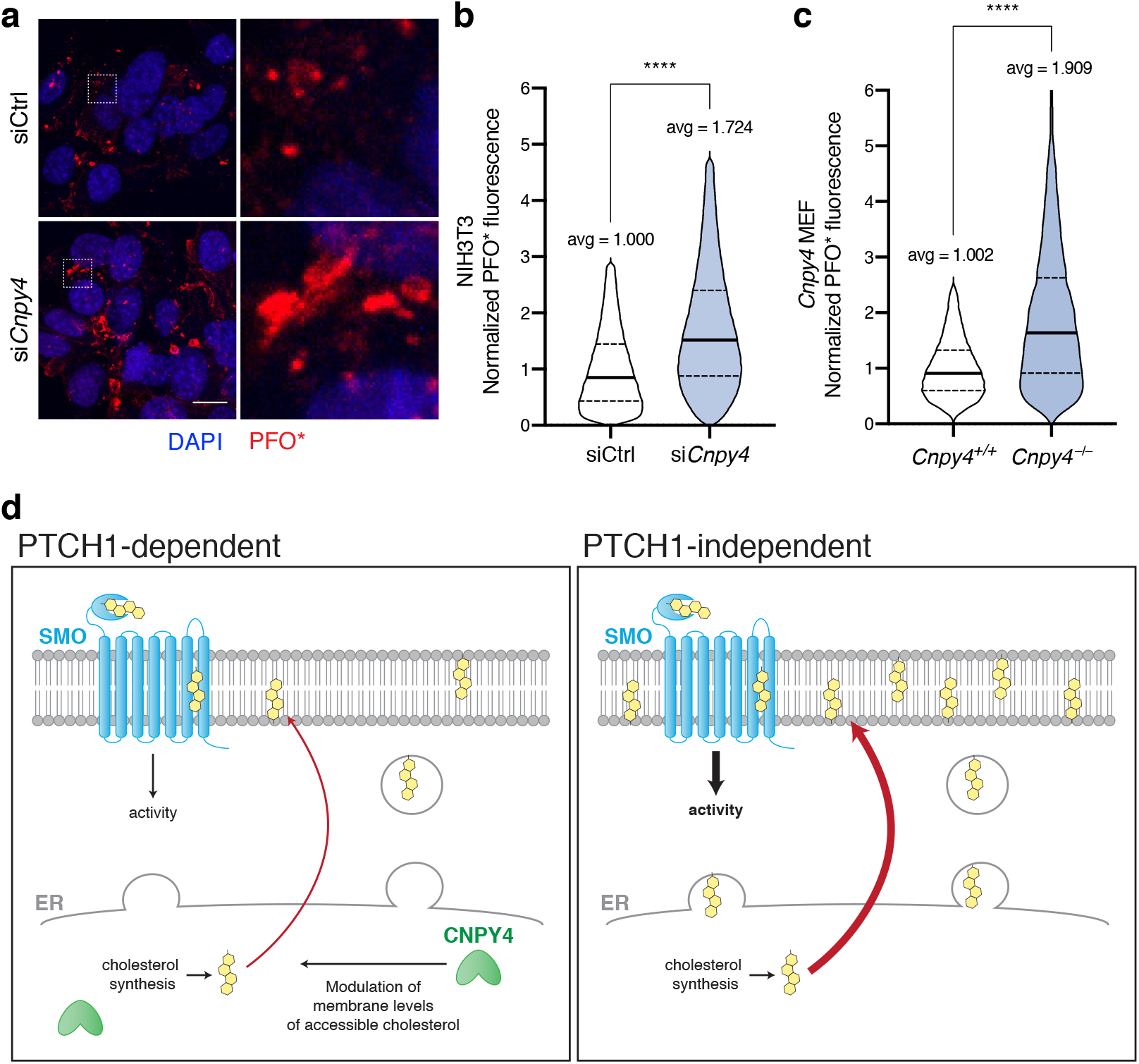
CNPY4 modulates levels of accessible cholesterol. **a**, Immunofluorescence of accessible cholesterol (PFO*-AF647, red) and nuclei (DAPI, blue) of NIH3T3 cells treated with *Cnpy4* or control siRNA. Boxed areas are magnified on the right. **b, c**, FACS analysis of NIH3T3 treated with *Cnpy4* or control siRNA (**b**) or *Cnpy4*^*+/+*^ and *Cnpy4*^−/−^ MEFs (**c**) stained with PFO*-AF647 for accessible cholesterol. Quantifications were normalized to the average value of control siRNA treated cells. Significance calculations were performed as described in Methods and Materials; **** p < 0.0001. **d**, Schematic illustrating CNPY4 modulation of HH activation. CNPY4, an ER-resident protein, likely modulates the ability of sterols, synthesized in the ER, to traffic to the ciliary membrane, thus modulating the ability of SMO to become activated in a manner parallel to that of PTCH1 regulation of SMO. In the absence of CNPY4, PTCH1 inhibition of SMO is bypassed by the elevated levels of membrane sterols, which lead to hyperactivation of HH signaling through SMO.

Taken together, our results reveal a new mechanism by which HH pathway activation is regulated via CNPY4-dependent control of sterol lipids in the membrane. Our data demonstrate that *Cnpy4*-dependent alteration of sterol lipid levels in the membrane directly modulate SMO-dependent HH activation. While increasing evidence supports the important role of lipids in signal transduction between PTCH1 and SMO, the molecular mechanisms governing these effects are not fully elucidated^29–35,41–45^. Recent studies have shown that both PTCH1 and SMO have several binding sites for sterols, including cholesterol, and that a subset of these binding events are essential for SMO activation^55,56,63–66^. Current models of HH activation propose that PTCH1 inhibits SMO by sequestering these activating sterols away from SMO^4,67^. This is thought to occur indirectly and to involve a proposed function of PTCH1 as a cholesterol pump, resulting in altered lipid composition of the plasma membrane^4,66^. Our data suggest that, similar in effect to PTCH1, but likely through a distinct, separate mechanism, CNPY4 regulates membrane composition to fine-tune SMO-dependent HH activation (Figure 4d). Our data also reveal that, by doing so, deletion of *Cnpy4* can bypass the PTCH1-inhibition of HH-activation (Figure 4d). Intriguingly, we additionally observed that depletion of *Cnpy4* causes a SMO-independent increase in basal HH activity, suggesting that accessible sterol levels in the membrane may contribute to HH signal transduction through multiple mechanisms. Recent work has illustrated that SMO movement into the cilia occurs even in the absence of ligand stimulation, albeit at much slower rates likely limited by diffusion^68,69^, and that sterol biosynthesis enzymes may regulate the ability of SMO to accumulate by priming the cilia via synthesis of sterols^52^. It is therefore possible that accessible sterol lipids affected by *Cnpy4* knockdown may be playing such a role, in addition to acting as a ligand for SMO activation.

How CNPY4 regulates the levels of accessible sterols at the plasma membrane remains an open question. CNPY4, as an ER-resident SAPLIP (Extended Data Fig. 8), is well positioned to assist with the synthesis, maturation, and membrane trafficking of lipids, such as sterols^70,71^. SAPLIP proteins can directly interact with lipids in the membrane in order to extract them for enzymatic presentation or membrane lysis^49–51^, and it possible that full-length CNPY4 maintains this functionality that we could not measure using the recombinant, truncated variant of CNPY4. Alternatively or possibly concurrently, CNPY4 might influence the maturation and membrane trafficking of components of the HH pathway, such as the SMO receptor. A similar function of CNPY4 was previously reported in the regulation of trafficking of Toll-like receptors^72,73^. Our findings elucidate a new regulatory modality in the HH pathway and identify a previously unknown function of CNPY4 as a regulator of HH pathway via modulation of plasma membrane sterols accessibility. This work provides a new context for unraveling the cellular mechanisms underlying previously reported functions of other CNPY proteins^49–51^.

## MATERIALS AND METHODS

### Mouse breeding

Mice were maintained in the University of California San Francisco (UCSF) specific pathogen-free animal facility in accordance with the guidelines established by the Institutional Animal Care and Use Committee and Laboratory Animal Resource Center. All experimental procedures were approved by the Laboratory Animal Resource Center at UCSF. Mice were maintained in temperature-controlled facilities with access to food and water *ad libitum*. *Cnpy4* heterozygote mice were produced by Lexicon (http://www.lexicon-genetics.com). To generate embryos at specific time points, adult mice were mated overnight, and females were checked for a vaginal plug in the morning. The presence of a vaginal plug was designated E0.5.

### Micro-computed tomography

Whole embryo or limb buds were collected and dehydrated through an ethanol series up to 70% ethanol. Samples were soaked in phosphotungstic acid (1%) overnight to differentially stain soft tissues as described previously^74^ and scanned using MicroXCT-200 (Carl Zeiss Microscopy) at 60 kV and 200 μA. We obtained 1200 projection images, taken at a total integration time of 3 seconds with linear magnification of 2x and a pixel size of 6.4 μm. The volume was reconstructed using a back projection filtered algorithm (Zeiss, Pleasanton, CA). Following reconstruction, tissues were manually segmented and rendered as 3-D surfaces using Avizo (FEI).

### Whole-mount *in situ* hybridization

Digoxygenin-labeled RNA probes (Roche) were generated by *in vitro* transcription from plasmids containing fragments of murine *Shh* and *Sox9*. Samples were fixed in 4% PFA overnight at 4°C, and the hybridization was carried out as previously described^9^.

### RNA-scope *in situ* hybridization

An RNAscope 2.5 HD Red (ACD, 310036, 322350) detection kit was used according to the manufacturer’s instructions. Sections were boiled in the target retrieval solution at ~100 °C for 15 min and incubated in the Protease Plus solution at 40 °C for 15 min. *Mus musculus Cnpy4* probe (475121 (lot # 16182A)) was used.

### Whole-mount *lacZ* staining

Embryos were fixed for 45 min in 4% PFA at 4°C, washed three times in rinse buffer containing 0.01% deoxycholate, 0.02% NP-40, 2 mM MgCl_2_, and 5 mM EGTA at room temperature and stained for 1 hr at 37°C in rinse buffer supplemented with 1 mg/mL X-gal, 5 mM K_3_Fe(CN)_6_, and 5 mM K_4_Fe(CN)_6_.

### Cell culture and drug treatments

Cells were cultured in Dulbecco’s modified Eagle media (Gibco) supplemented with 10% FBS (Hyclone) and penicillin streptomycin (Gibco) and incubated at 3°7C with 5% CO_2_. All cells lines were regularly tested for mycoplasma contamination using the MycoAlert mycoplasma detection kit (Lonza). Stimulations were performed in low-serum OptiMEM (Life Technologies) to induce ciliation with 100 nM SAG (EMD Millipore), 1 μg/mL recombinant SHH (R&D Systems), 30 μM 20(S)-hydroxycholesterol (Cayman Chemicals), 30 μM 24(S), 25-epoxycholesterol (Avanti Polar Lipids), or 25 μM SANT-1 (Selleckchem). Incubations with SAG, SHH, and SANT-1 were done for 24 hours and oxysterols were done for 30-36 hours.

### MEF generation

Embryos were isolated and washed in 1x PBS twice. Limb buds were separated using sterile tweezers from each embryo and washed with DMEM before incubation with 0.25% Trypsin/EDTA (Gibco) at 37°C for 10 minutes. Trypsin was quenched by addition of DMEM supplemented with FBS and penicillin streptomycin. Cells were pipetted up and down at least 10 times to further dissociate cells before being transferred into fresh 15 mL tubes. Cells were gently pelleted at 200xg for 5 minutes at room temperature. Supernatant was carefully aspirated and cells were resuspended in fresh media and plated in 6-cm plates (Gibco). Additional cell debris was aspirated off and fresh media added daily until cells reached confluency, upon which they were split and expanded once before being pooled and flash frozen.

### siRNA transfection

22.5 pmol of siRNA SMARTpool (Dharmacon) were transiently transfected into indicated cells using lipofectamine RNAiMax (Invitrogen) according to the manufacturer’s protocols. Cells were transfected for 72 hours before cell analysis. Confirmation of mRNA silencing was done by qRT-PCR analysis and confirmation of protein knockdown was performed via Western blotting.

### qRT-PCR analysis

Cells were grown in either 6- or 12-well plates and treated with indicated expression conditions. RNA was extracted from cells using the RNEasy Mini kit (Qiagen) and reverse-transcribed to produce cDNA using the iScript cDNA synthesis kit (Bio-Rad). qRT-PCR was performed using Power-Up SYBR Green Master Mix (Applied Biosystems) on an Invitrogen real-time PCR machine. mRNA transcript relative abundances were calculated using the ΔΔCt method against *Gapdh*.

**Table 1.1.**
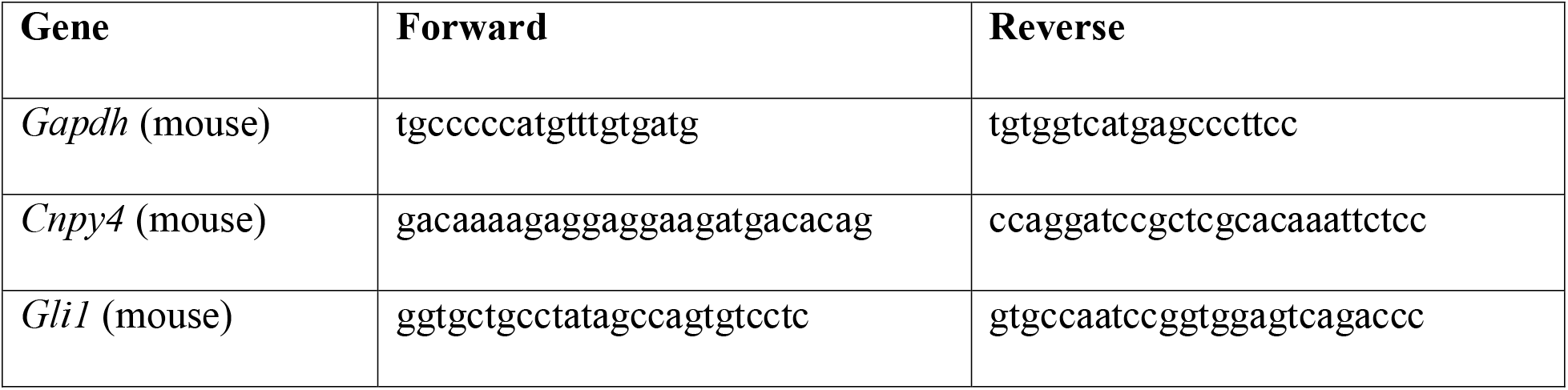
qRT-PCR primers.

### Luciferase-based reporter assays

Cells were plated in 6-well plates and transfected with siRNA as described above at least 16 hours post-plating. 396 ng of Gli1-responsive Firefly luciferase reporter plasmid, 4 ng of a control Renilla luciferase reporter plasmid under the control of a constitutively active TK promoter, and 1 μg of pcDNA3.1+ empty vector were transfected into cells at least 6 hours post-siRNA transfection using lipofectamine LTX with Plus reagent (Invitrogen) according to the manufacturer’s protocol. 16 hours-post transfection, cells were recovered with fresh media for 24 hours. Stimulation with indicated ligand was performed in low-serum OptiMEM media (Gibco) for 24-36 hours. Luciferase assays were conducted using the Dual Luciferase Reporter Assay System (Promega) and measured on a GloMax 96 Microplate Luminometer with Dual Injectors (Promega).

### Immunofluorescence staining

NIH3T3 or COS-7 cells were plated onto glass coverslips and transfected the following day. Cells were fixed in 3.7% PFA solution diluted in 1x PBS at room temperature with rocking and then incubated with a 0.1% Triton X-100 and 2.5% BSA solution in 1x PBS to permeabilize cells and to block for non-specific antibody interaction. Primary antibodies were diluted in blocking buffer and incubated overnight at 4°C then washed out three times with 0.1% Triton X-100 in 1x PBS. Secondary antibodies were diluted in blocking buffer and incubated for 2 hours at root temperatures before subsequent washes. DAPI staining was conducted for 10 minutes following the last wash before cells were mounted onto glass coverslips with Prolong Gold AntiFade Mountant (Life Technologies).

### PFO* staining and FACS analysis

Cells were grown in 6-wells and treated with indicated conditions. Cells were lifted with 0.5% Triton-EDTA and gently pelleted by centrifugation at 200xg for 5 minutes. Pellets were washed gently two times with 1x PBS before incubation in blocking buffer (10 mg/mL BSA in 1x PBS) for 10 minutes on ice. Cells were pelleted once more before incubation with 5 μg/mL PFO* probe diluted in blocking buffer for 30 minutes on ice. Cells were gently washed one time with 1x PBS before analysis by FACS. Fluorescent intensity measurements by flow cytometry were performed on a Sony Cell Sorter SH800 using a 638 nm laser for excitation. Live and singlet populations were selected based on forward and side scatter. No further gating was used to select cell populations. Outliers were identified using the identify outliers function on Prism 8 (GraphPad).

### Microscopy

Bright-field images were acquired on an Axio Imager.Z2 upright microscope (ZEISS) for whole mount, *in situ* hybridization and *lacZ* staining. Immunofluorescence and PFO* images were acquired on either a Nikon Elipse Ti with a CSU-X1 spinning disc confocal and Andor Clara interline CCD camera with a Nikon Plan Apo 60x oil objective or a Zeiss LSM 800 confocal laser scanning microscope with a 63x oil objective. Cell length calculations and SMO intensity analysis was done on Fiji.

### Recombinant protein expression and purification

CNPY4 constructs were synthesized by Genscript and subcloned into a pET28b plasmids with a 10xHis tag sequence. Cloning verification was done by DNA sequencing (Elim biotechnology). Constructs were transformed into SHuffle T7 competent *E. coli* cells (NEB) and underwent antibiotic selection on Kanamycin plates for 16 hours at 37°C. A single colony was used to inoculate a Luria broth starter culture supplemented with appropriate antibiotic for 16 hours at 37°C and 220 rpm shaking. 10 mL of starter culture was used to inoculate 900 mL of Terrific broth supplemented with appropriate antibiotic and 100 mL of 10x phosphate buffer (0.17 M KH_2_PO_4_, 0.72 M K_2_HPO_4_). Cells were grown at 37°C, 220 rpm shaking to an OD_600_ of 0.6 - 0.8 before being induced with isopropyl β-D-1-thiogalactopyranoside to a final concentration of 0.5 mM. Cultures were grown for an additional 20 hours at 18°C and 220 rpm shaking. Cells were collected by centrifugation using an Avanti centrifuge equipped with a JA 8.5 rotor at 4000xg, 40 minutes, 4°C. Pellets were flash frozen for later purification or resuspended in binding buffer (50 mM HEPES, pH 8.0, 500 mM NaCl, 20 mM imidazole, pH 8.0, 5% glycerol) supplemented with DNaseI (Sigma Aldrich) and cOmplete mini EDTA-free protease inhibitor cocktail (Roche) and lysed via sonication at 30% amplitude, 4 seconds on, 2 seconds off, for a total of 5 minutes. Lysates were clarified in an Avanti centrifuge equipped with a JLA 25.50 rotor at 20,000xg, 40 minutes, 4°C. Clarified lysates were incubated with Ni-NTA 6 Fast Flow beads (GE Life Sciences), with gentle rotation, for 16 hours at 4°C before being applied to a gravity flow Econo-column (Bio-Rad). Beads were washed thoroughly with 20 column volumes of binding buffer followed by 10 column volumes of binding buffer supplemented with an additional 12.5 mM imidazole. The recombinant protien was eluted in 5 column volumes of elution buffer (binding buffer with 250 mM imidazole). The elution was buffer exchanged back into low Imidazole binding buffer and incubated with 1 mg of recombinant 3C protease for 16 hours at 4°C. Uncleaved protein was removed by passing over fresh Ni-NTA 6 resin. The protein was then diluted 10 times with mono Q binding buffer (50 mM HEPES, pH 8.0) and applied to a MonoQ 5/50 GL column (GE Life Sciences) connected to an AKTA Pure system (GE Life Sciences).. Recombinant hCNPY4ΔCt was eluted with a linear gradient of elution buffer (50 mM HEPES, pH 8.0, 500 mM NaCl). Elutions were concentrated using an Amicon Ultra-15 10k MWCO centrifugal filter (Millipore) before being loaded onto a a Supderdex 200 16/600 column (GE Life Sciences) equilibrated in size exclusion chromatography (SEC) buffer (50 mM Bicine, pH 9.0, 150 mM NaCl). Fractions confirmed to contain pure hCNPY4ΔCt by SDS-PAGE analysis were pooled and flash-frozen in liquid nitrogen and stored at −80°C.

### Circular dichroism

Purified CNPY proteins were analyzed on a Jasco J-810 spectropolarimeter at 1 nm steps. Proteins were analyzed at an approximate concentration of 2 μM in a 50 mM sodium phosphate buffer, pH7.0 at 25°C. Thermal melt data was collected at 222 nm with a temperatures range of 25°C to 95°C in increments of 5°C. CNPY4ΔCt was additionally incubated with 30 μM of cholesterol, 20(S)-hydroxycholesterol, or 24(S), 25-epoxycholesterol prior to thermal melt analysis for assessment of binding capacity. Data for three, averaged reads was fitted using the log(agonist) vs. response -- Variable slope non-linear analysis on Prism 8 (GraphPad), and the LogEC50 from the analysis was reported as the melting temperature. Error bars correspond to calculated standard error of the mean.

### Fluorescence polarization

Purified human CNPY4ΔCt in SEC buffer (50 mM HEPES, pH 8.0, 150 mM NaCl) were analyzed for binding to 50 nM BODIPY-cholesterol (Cayman Chemical) at the indicated protein concentrations. 1% Tween-20 was added to the reaction mixture. Experiments were performed with a reaction volume of 20 μL in triplicate using a black-bottom 384-well plates (Corning) on an Analyst AD plate reader (Molecular Devices). Excitation and emission wavelengths used for the kinetic experiments were 480 nm and 508 nm, respectively, according to the manufacturer’s recommendation. Kinetic reads were performed over 15 minutes with 30 second intervals. As no significant difference was observed, signal was averaged across all time points for each triplicate with standard error calculated for each data point. Data was fitting using the Semilog line -- X is log, Y is linear non-linear analysis on Prism 8 (GraphPad). Error bars correspond to calculated standard error of the mean.

### Statistical analysis

All statistical analysis were performed using Prism 8 (GraphPad). Significance analysis for luciferase assay (N=9; 3 biologic replicates and 3 technical replicates) and qRT-PCR (N=12; 4 biologic replicates and 4 technical replicates) analyses were done using the Mann-Whitney non-parametric test. Ciliation (N=77 with t=1.785, df=152 for NIH3T3 cells; N=79 with t=1.855, df=156 for MEF cells) and FACS analyses (N=92183 for *Cnpy4*^*+/+*^ MEFs and N=74848 for *Cnpy4^−/−^* MEFs with t=183.1 and df=95074; N=244550 for siCtrl NIH3T3 cells and N=291375 for si*Cnpy4* NIH3T3 cells with t=295.6, df=503909) were performed using the Welch’s t-test. All statistical analyses were two-tailed.

## Supporting information

Supplemental figures all

## ACKNOWLEDGMENTS

We thank Pao-Tien Chuang and Jeremey Reiter for the NIH3T3, *Sufu*^−/−^, and *Ptch1*^−/−^ MEF cell lines, Maia Kinnebrew and Rajat Rohatgi for the PFO* probe, Shao-qing Zhang and Haifan Wu for their guidance on CD experiments, Vikas Daggubati for assistance with experimental procedures, and Sarah Findakly-Oshima for help with confocal imaging. We also thank Tom Kornberg, Aashish Manglik, Kara McKinley, and Jennifer Kung for critical reading of the text and members of the Jura and Klein labs for their helpful discussions. This work was funded by the UCSF Program for Breakthrough Biomedical Research and NIDCR R01-DE028496 and R35-DE026602 to O.D.K.

## AUTHOR CONTRIBUTIONS

N.J. and O.D.K. conceived of and initiated the project. M.L. performed the in cell-based experiments, recombinant protein expression and purifcation, and *in vitro* experiments. A.S. generated the mouse models and performed the *in situ* hybridizations. M.D.P, H.T., and C.A. assisted with the *in vitro* experiments. M.L., N.J., and O.D.K. wrote the manuscript with provisions provided by D.R.R.

