## Supplemental figures all for "CNPY4 inhibits the Hedgehog pathway by modulating membrane sterol lipids"

Extended Data Figure 1, Lo et al.

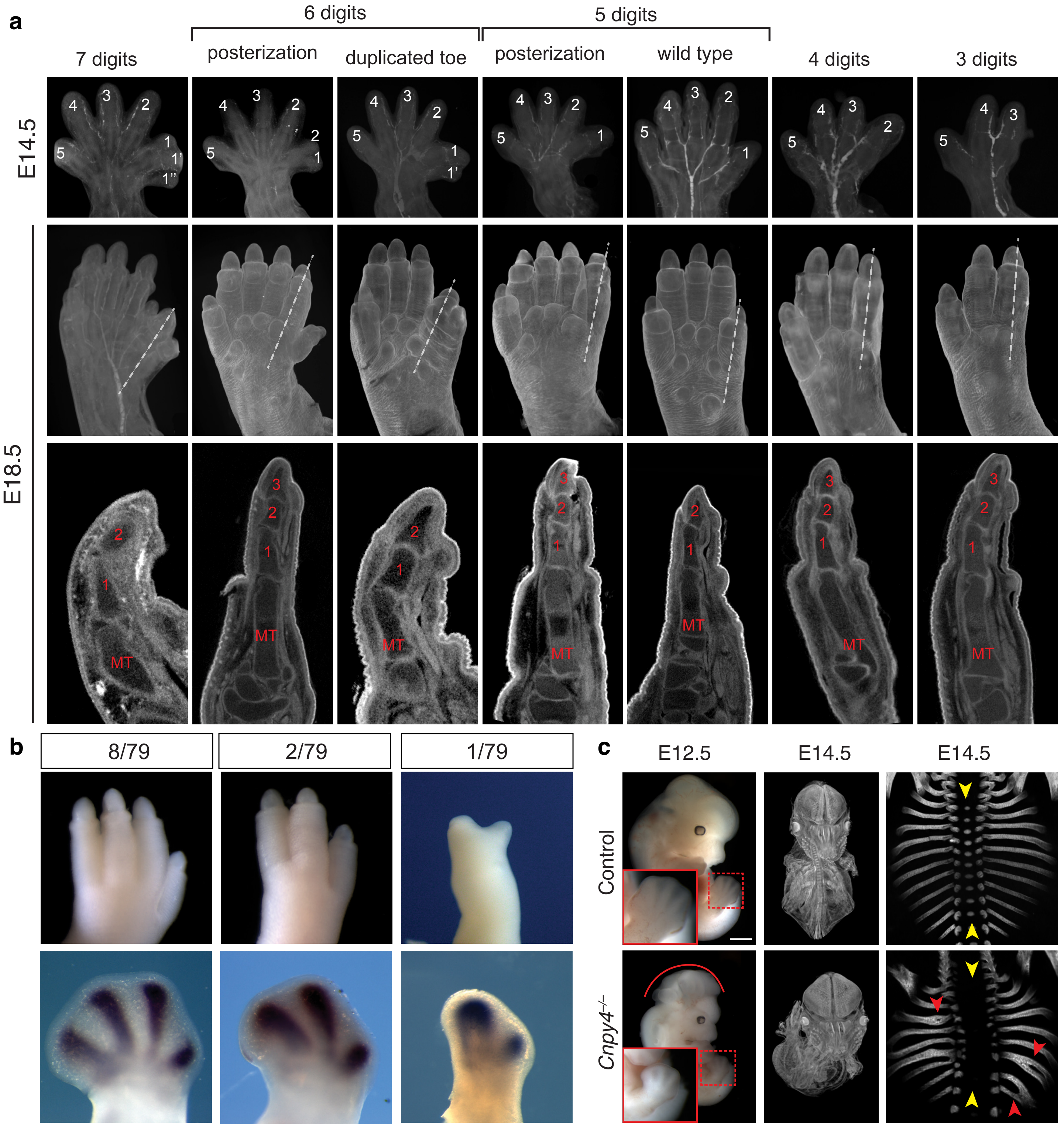

Extended Data Figure 2, Lo et al.

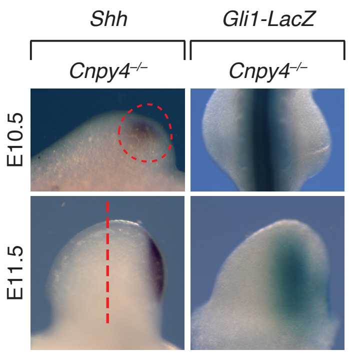

Extended Data Figure 3, Lo et al.

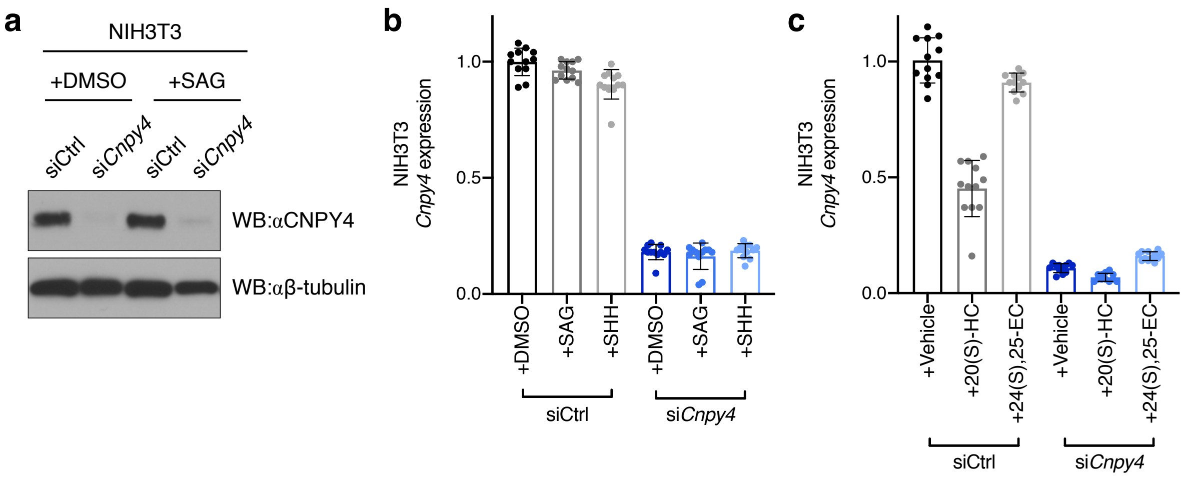

Extended Data Figure 4, Lo et al.

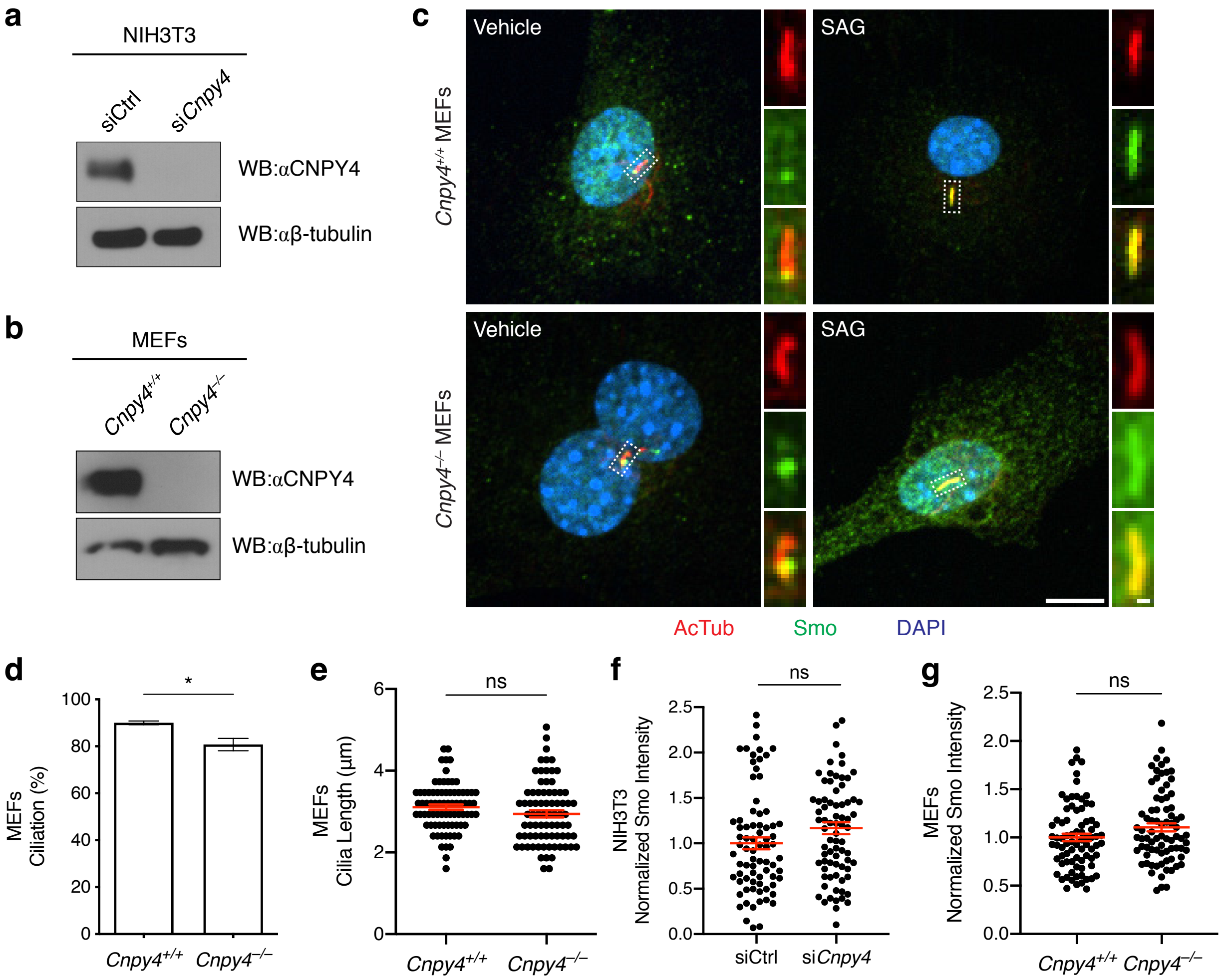

Extended Data Figure 5, Lo et al.

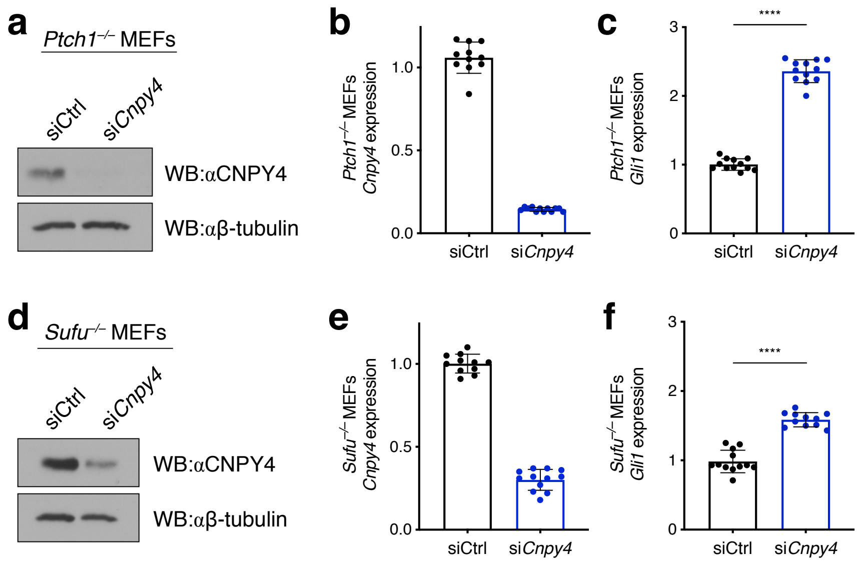

Extended Data Figure 6, Lo et al.

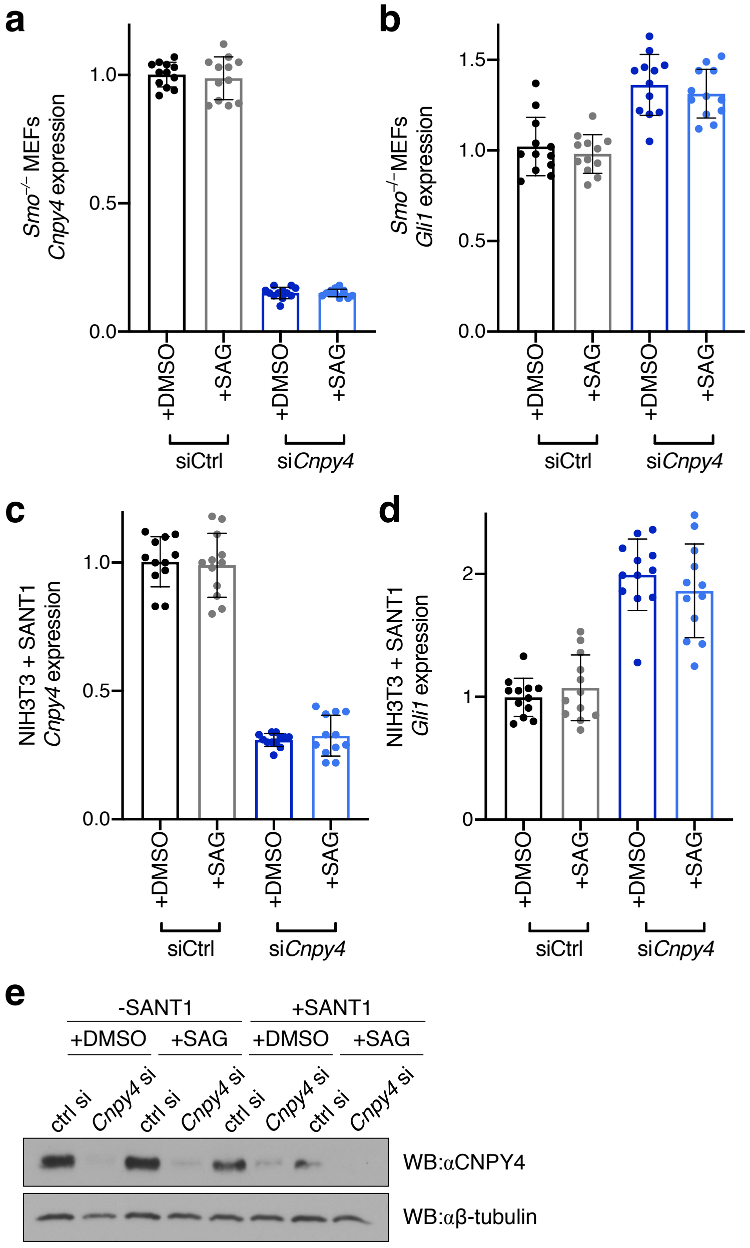

Extended Data Figure 7, Lo et al.

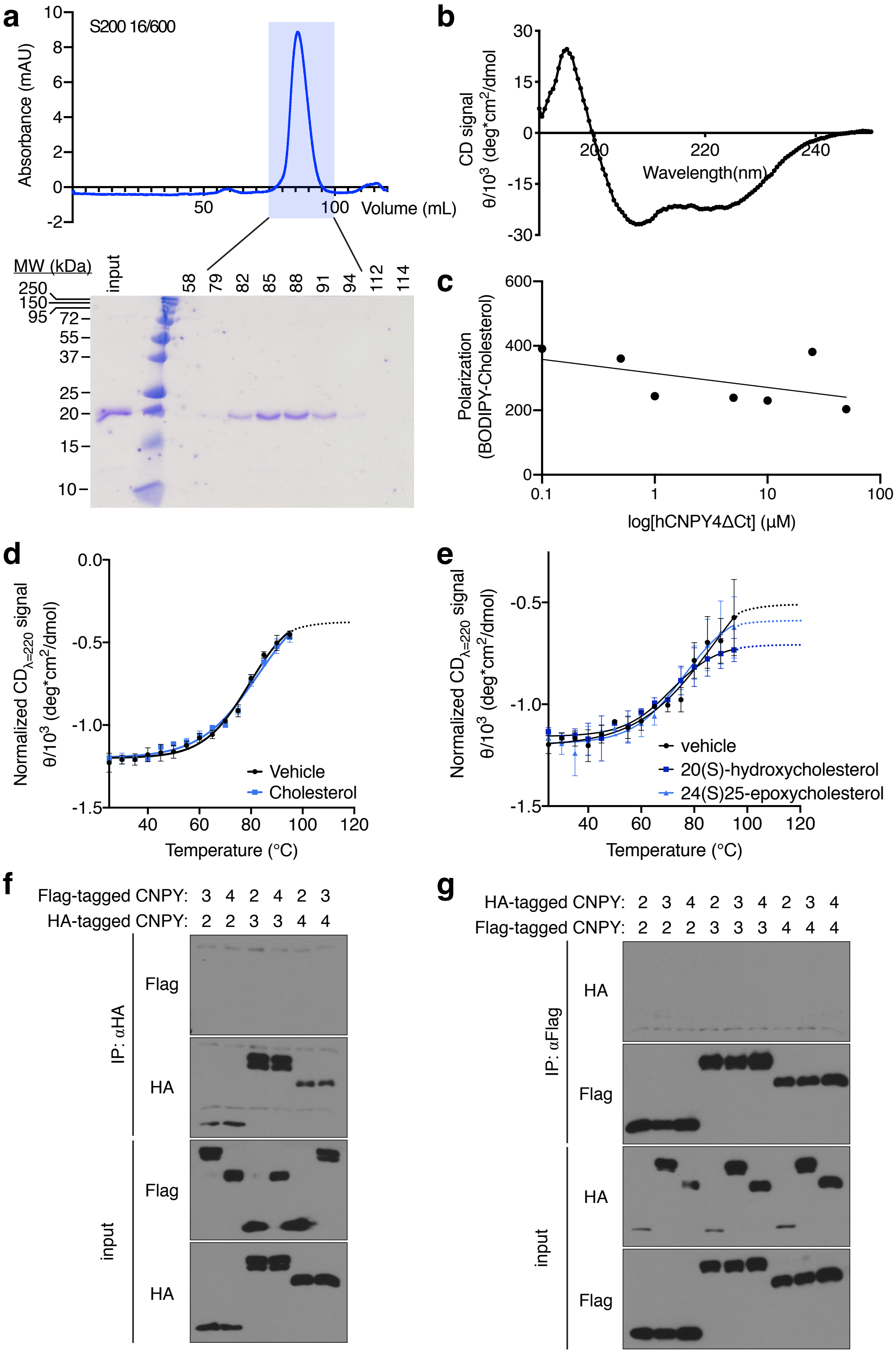

Extended Data Figure 8, Lo et al.

**a**

```
SAP_B        1---------------------------------------------NGDVCQDCIQMVTDIQTAVRTNSTF-------------------------
CNPY1        1 -----------------------------------------------------------------------------------------------
CNPY2        1 -------------MKGWGWLALLLGALLGTAWAR---------RSQDLHCGACRALVDELEWEIAQVDPKKTIQMGSFRINPDGSQSVVEVPYAR
CNPY3        1 MDSMPEPASRCLLLLPLLLLLLLLL-PAPELGPSQAGAEENDWVRLPSKCEVCKYVAVELKSAFEETGKTKEVIGTGY--GILD-QKASGVKYTK
CNPY4        1 -----------MGPVRLGILLFLFL-AVHEAWAGMLKEEDDDTERLPSKCEVCKLLSTELQAELSRTGRSREVLELGQ--VLDTGKRKRHVPYSV
consensus    1                    .  .                         ..  .. .  ..   .      . .                 . .

SAP_B       26 -VQALVEHVKEECDRLGP---------------------------------------------------GMADICKNYISQYSEIAIQMMMHMQP
CNPY1        1 ---------------MNDYKLEEDPVTKERTFKRFA---------PRKGDKIYQEFKK--LYFYSDAYRPLKFACETIIEEYEDEISSLIAQE-T
CNPY2       74 SEAHLTELLEEICDRMKEYGEQIDPSTHRKNYVRVV---------GRNGESSELDLQG--IRIDSDISGTLKFACESIVEEYEDELIEFFSRE-A
CNPY3       92 SDLRLIEVTETICKRLLDYSLHKERTGSNRFAKGMSETFETLHNLVHKGVKVVMDIPYELWNETSAEVADLKKQCDVLVEEFEEVIEDWYRNHQE
CNPY4       82 SETRLEEALENLCERILDYSVHAERKGSLRYAKGQSQTMATLKGLVQKGVKVDLGIPLELWDEPSVEVTYLKKQCETMLEEFEDIVGDWYFHHQE
consensus   96 .. .. . .. ..... .. .  .     .  .             ... .. ...        .     ..  *........... ... . .

SAP_B       69 KEI----CA-LVGFCDEVK----------------------------------------------------------------------------
CNPY1       69 HYLADKLCSEKSDLCETSANHTE-----------L------------------------------------------------------------
CNPY2      157 DNVKDKLCSKRTDLCDHALHISH---------DEL------------------------------------------------------------
CNPY3      187 EDLTEFLCANHVLKGKDTSCLAEQWSGKKGDTAAL---GGKKSKKKSSRAKAAGGRSSSSKQRKELGGLEGDPSPEEDEGIQKASPLTHSPPDEL
CNPY4      177 QPLQNFLCEGHVLPAAETACLQETWTGKEITDGEEKTEGEEEQEEEEEEEEEEGGDKMTKT--------GSHPKLD-RE--------------DL
consensus  191   . . .*  ..  ....  . .           .
```

**b**

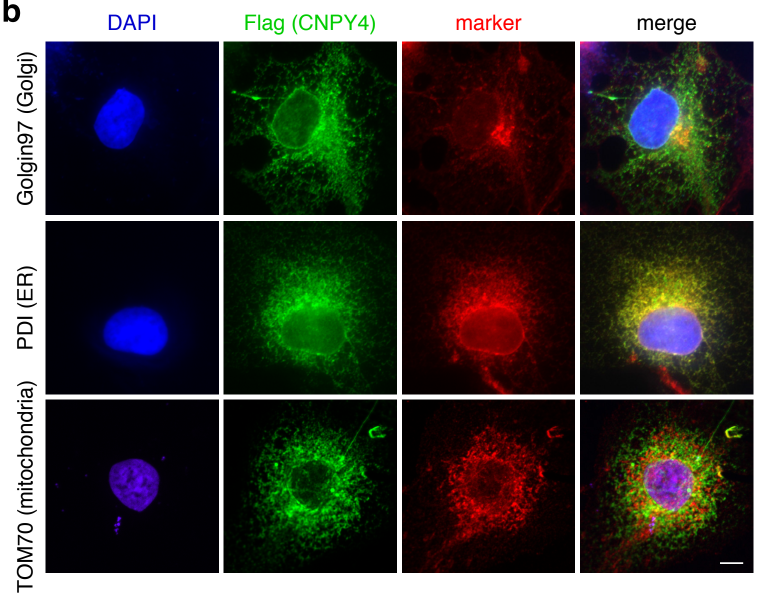

**Extended Data Fig. 1 | Developmental defects in a *Cnpy4* knockout mouse model. a,** Micro computed tomography (μCT) reconstructions of *Cnpy4* mutant limbs at E14.5 (top row) and E18.5 (bottom two rows), showing a range of hindlimb digit phenotypes from oligodactyly to polydactyly. **a'** sagittal plan images through digit 1 (dash line in middle row) showing the number of phalanges (bottom). **b**, Dorsal view of *Cnpy4* mutant limbs at E18.5 (top) and whole mount in situ hybridization for *Sox9* at E12.5 (bottom) showing oligodactyly phenotypes. **c**, Whole mount images and μCT analysis showing (left) cranial neural tube defect (exencephaly); (middle) shortening and kinking of the body axis; (right) abnormal rib morphology, with fusions and bifurcations and truncation of the sternum.

**Extended Data Fig. 2 | Reduction of *Shh* expression in minority of *Cnpy4* mutants.** *In situ* hybridization of *Shh* (left) and *Gli1-LacZ* (right) in hindlimb buds at E10.5 and E11.5 showing reduced expression in a minority of *Cnpy4* mutants.

**Extended Data Fig. 3 | Knockdown of *Cnpy4* in NIH3T3 cells. a**, Western blot analysis of CNPY4 and β-tubulin loading control proteins levels of NIH3T3 cells treated *Cnpy4* or control siRNA. **b, c**, qRT-PCR assessment of *Cnpy4* expression in ciliated NIH3T3 cells treated with *Cnpy4* or control siRNA and stimulated with SAG or recombinant SHH (**b**) and 20(S)-hydroxycholesterol or 24(S), 25-epoxycholesterol (**c**).

**Extended Data Fig. 4 | Loss of CNPY4 has little effect on the primary cilia. a, b**, Western blot analysis of CNPY4 and β-tubulin loading control proteins levels of NIH3T3 cells treated *Cnpy4* or control siRNA (**a**) or control and *Cnpy4* null MEFs (**b**) used for immunofluorescence. **c**, Immunofluorescence of primary cilia (acetylated tubulin, red), SMO (SMO, green), and the nuclei (DAPI, blue) in control and *Cnpy4* null MEFs with SAG or vehicle (DMSO) treatment. The scale bar represents 10 um. Inset scale bar represents 1 um. **d**, Quantification of number of MEFs ciliated as assessed by acetylated tubulin immunofluorescence. **e**, Quantification of ciliary length in MEFs. Measurements were performed in FIJI using the acetylated tubulin channel. **f, g**, Quantification of SMO trafficking to cilia in NIH3T3 (**f**) and MEF (**g**) cells. Analyses were done using FIJI by measuring the average fluoresence intensity of SMO over the length of the cilia in the appropriate channel. Average background fluorescence measured over the same length from NIH3T3 cell images were subtracted. Average background fluorescence in MEF cell images deviated by <5% and were not subtracted. All significance calculations were performed as described in Methods and Materials; ns = $p > 0.05$, * $p < 0.05$.

**Extended Data Fig. 5 | *Cnpy4* epistatic interaction with HH pathway components. a**, Western blot analysis of CNPY4 and β-tubulin loading control proteins levels in *Ptch1*$^{-/-}$ MEF cells with *Cnpy4* or control siRNA treatment. **b, c**, qRT-PCR assessment of *Cnpy4* (**b**) and *Gli1* (**c**) expression in *Ptch1* null MEFs. **d**, Western blot analysis of CNPY4 and β-tubulin loading control proteins levels in *Sufu*$^{-/-}$ MEF cells with *Cnpy4* or control siRNA treatment. **e, f**, qRT-PCR assessment of *Cnpy4* (**e**) and *Gli1* (**f**) expression in *Sufu* null MEFs. All significance calculations were performed as described in Methods and Materials; **** $p < 0.0001$.

**Extended Data Fig. 6 | SHH signal regulation by CNPY4 requires SMO. a, b**, qRT-PCR assessment of *Cnpy4* (**a**) or *Gli1* (**b**) expression in ciliated *Smo*$^{-/-}$ MEF cells treated with *Cnpy4* or control siRNA and stimulated with SAG. **c–d**, qRT-PCR assessment of *Cnpy4* (**c**) or *Gli1* (**d**) expression in ciliated NIH3T3 cells treated with SANT-1 and *Cnpy4* or control siRNA before stimulation with SAG. **e**, Western blot analysis of CNPY4 and β-tubulin loading control proteins levels of NIH3T3 cells treated *Cnpy4* or control siRNA, SANT-1, and SAG stimulation.

**Extended Data Fig. 7 | Recombinant CNPY4 does not bind oxysterols involved in SHH activation. a**, Size exclusion chromatography profile for hCNPY4ΔCt purification with accompanying gel stained with Coomassie. **b**, Circular dichroism of hCNPY4ΔCt at room temperature**. c**, Fluorescence polarization of a BODIPY-cholesterol probe incubated with increasing concentration of purified hCNPY4ΔCt. **d**, Thermal melt of hCNPY4ΔCt incubated with vehicle (chloroform) or cholesterol analyzed by circular dichroism at a wavelength of 222 nm. **e**, Thermal melt of hCNPY4ΔCt incubated with vehicle (methyl-β-cyclodextrin), 20(S)-hydroxycholesterol, or 24(S), 25-epoxycholesterol analyzed by circular dichroism at a wavelength of 222 nm. **f, g**, Co-immunoprecipitation or Flag-tagged and HA-tagged variants of wild-type CNPY2, CNPY3, and CNPY4 for homo- and hetero-dimerization. Proteins were transiently expressed in HEK293 cells, pulled-down using either an anti-HA (**f**) or an anti-Flag (**g**) antibody. Protein levels were detected with the indicated antibodies by Western blot analysis.

**Extended Data Fig. 8 | CNPY4 is an ER-resident member of the SAPLIP family. a**, Protein sequence alignment for human Saposin B, CNPY1, CNPY2, CNPY3, and CNPY4. Conservation was assigned using SIM-Alignment tool (ExPASy). **b**, Immunofluorescence of COS-7 cells transiently co-transfected with Flag-tagged human CNPY4. CNPY4 was detected with an anti-Flag antibody (green), the nucleus with DAPI (blue), and markers for organelles with the indicated antibody (red). Scale bar corresponds to 10 μm.
